# TIR signaling activates caspase-like immunity in bacteria

**DOI:** 10.1101/2024.10.24.620036

**Authors:** François Rousset, Ilya Osterman, Tali Scherf, Alla H. Falkovich, Azita Leavitt, Gil Amitai, Sapir Shir, Sergey Malitsky, Maxim Itkin, Alon Savidor, Rotem Sorek

## Abstract

Proteases of the caspase family, as well as Toll/Interleukin-1 Receptor (TIR)-domain proteins, have central roles in innate immunity and regulated cell death in humans. In this study we describe a bacterial immune system comprising both a caspase-like protease and a TIR-domain protein. We found that the TIR protein, once it recognizes phage invasion, produces the previously unknown immune signaling molecule ADP-cyclo[N7:1′′]-ribose (N7-cADPR). This molecule specifically activates the bacterial caspase-like protease which then indiscriminately degrades cellular proteins to halt phage replication. The TIR-caspase defense system, which we denote as type IV Thoeris, is abundant in bacteria and efficiently protects against phage propagation. Our study highlights the diversity of TIR-produced immune signaling molecules and demonstrates that cell death regulated by proteases of the caspase family is an ancient mechanism of innate immunity.

## Introduction

Caspases are proteases with central roles in innate immunity and regulated cell death in humans^1^. The human genome harbors twelve caspase-encoding genes, with some of them promoting inflammation while others functioning in the execution of cell death^1^. Inflammatory caspases, including caspases 1, 4 and 5, are recruited to activated inflammasomes following pathogen recognition, and are responsible for the proteolytic activation of cytokines and gasdermin D, promoting cell death through pyroptosis^2^. Executioner caspases (e.g., caspases 3, 6 and 7), once activated by initiator caspases (8, 9 and 10), cleave hundreds of protein substrates to promote apoptotic cell death^1,3^. Proteases of the caspase family (Pfam PF00656) also exist in prokaryotes^4^, and were proposed to cleave bacterial gasdermins to activate immunity in response to phage infection^5^. Bacterial proteins of the caspase family were also shown to promote cell death when activated by type III CRISPR-Cas systems^6–8^. However, the molecular functions of the vast majority of caspase-like proteins in bacteria remain poorly understood^4^.

Toll/Interleukin-1 Receptor (TIR) domain proteins also play key roles in innate immunity. Initially described as protein-protein interaction modules in human Toll-like and interleukin-1 receptors^9^, TIR domains in bacteria and plants were later shown to be enzymes that produce immune signaling molecules using nicotinamide adenine dinucleotide (NAD^+^) as a substrate^10^. In the bacterial defense system type I Thoeris, the TIR-domain protein, once it senses phage infection, produces the signaling molecule 1”-3’ glycocyclic ADP-ribose (gcADPR)^11,12^. This molecule activates an effector protein that depletes the cell of NAD^+^ and aborts phage replication^13,14^. The TIR-domain protein in type II Thoeris, on the other hand, produces the signaling molecule histidine-ADP-ribose (His-ADPR)^15^, which activates an effector protein that disrupts the cell membrane to abort phage propagation. A third type of Thoeris was recently proposed but its mechanism of action and molecular signaling remain unknown^16^. In plants, TIR-derived signaling molecules are diverse and include phosphoribosyl-AMP/ADP^17^, 1”-2’ gcADPR^11,12,18^, di-ADPR^19^, ATP-ADPR^19^ and 2’,3’-cAMP/cGMP^20^. Current evidence suggests that the repertoire of immune signaling molecules produced by TIR domains has not been fully unveiled^21^.

In this study, we describe type IV Thoeris, a bacterial defense system encoding both a TIR-domain protein and a caspase-like protease. We show that upon phage infection, the TIR-domain protein produces a novel signaling molecule, N7-cADPR, that binds the caspase-like protease and triggers promiscuous arginine-specific protease activity. Protease activation leads to indiscriminate cleavage of multiple proteins, including elongation factor Tu (EF-Tu), thereby aborting phage infection. Our study establishes a direct functional connection between TIR signaling and caspase activation in bacterial defense against phages.

## Results

### A TIR-caspase operon provides anti-phage immunity

While examining the genomic environment of caspase-like proteins in bacteria, we noticed an abundant two-gene operon encoding a short TIR-domain protein and a caspase-like protease (**Fig. 1A**). Operons with this gene organization were frequently encoded in the vicinity of bacterial defense systems, suggesting a defensive function (**Fig. S1**). We synthesized and cloned two such operons, one from *Escherichia coli* 328 and the other from *Pseudomonas sp.* 1-7, and expressed these in *E. coli* K-12 MG1655 under the control of an arabinose-inducible promoter. Following a challenge by a panel of phages, we observed that the infectivity of phage T6 was substantially reduced when plated on cells expressing either of the operons, showing that this TIR- and caspase-encoding operon is a novel defense system (**Fig. 1B**). Since the *E. coli* 328 operon exhibited slight toxicity upon expression induction, we selected the *Pseudomonas sp.* 1-7 (*Ps*) homolog for further study.

**Figure 1.**
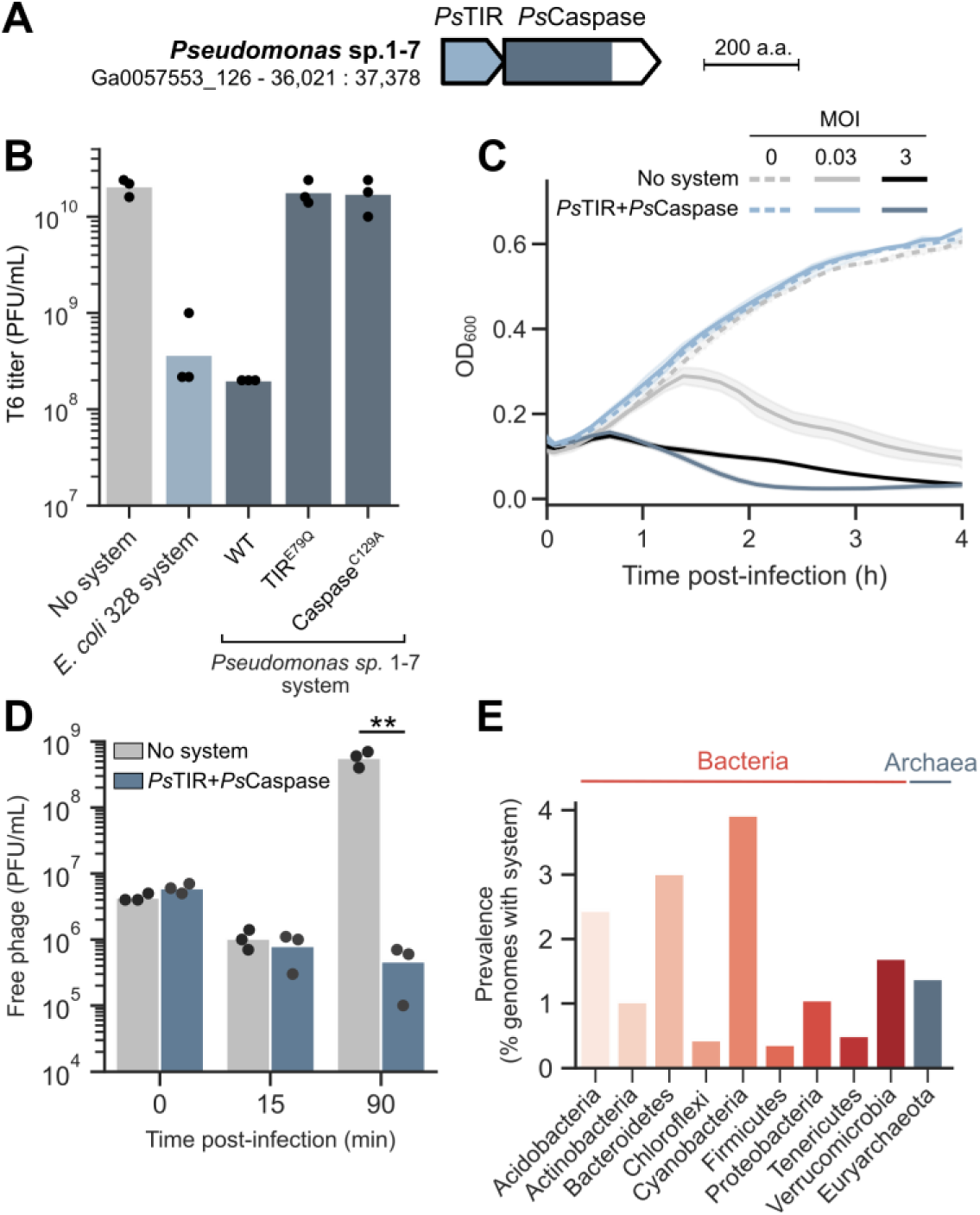
A genetic system encoding a TIR-domain protein and a caspase-like protease provides anti-phage defense. (**A**) Genetic architecture of a system from *Pseudomonas* sp. 1-7. IMG^23^ genome ID and gene coordinates are displayed below strain name. TIR and caspase-like domains are shown as blue shades. (**B**) Quantification of phage infection efficiency via plaque assays. Tenfold serial dilutions of phage T6 were spotted on a lawn of *E. coli* K-12 MG1655 cells expressing an empty vector (no system), the system from *E. coli* 328, or the system from *Pseudomonas* sp. 1-7, either wild-type or mutated in the predicted active sites of the TIR (E79Q) or caspase-like protease (C129A). Bars show the mean of three replicates with individual data points overlaid. (**C**) Growth curves of *E. coli* K-12 MG1655 cells expressing an empty vector or the system from *Pseudomonas* sp. 1-7, infected by phage T6 at an MOI of 0.03 or 3 (or 0 for uninfected cells). Curves show the mean of three replicates with the standard deviation shown as a shaded area. (**D**) Plaque-forming units of phage T6 sampled from the supernatant of *E. coli* K-12 MG1655 cells expressing an empty vector or the system from *Pseudomonas* sp. 1-7. Cells were infected at an MOI of 0.1. Bars represent the mean of three replicates with individual data points overlaid. Stars show significance of a two-sided t-test (**p < 0.01). (**E**) Detection of the defense system in prokaryotic genomes, shown for phyla with at least 50 genomes and where at least one system was detected.

Single amino acid substitutions in the predicted catalytic sites of *Ps*TIR (E79Q) and *Ps*Caspase (C129A) proteins abolished defense (**Fig. 1B**), suggesting that the enzymatic activity of both proteins is required for immunity. Bacterial cells expressing the system were able to survive when infected with T6 in liquid culture at a low multiplicity of infection (MOI), but died when infected at a high MOI (**Fig. 1C**). In addition, infected cells expressing the system did not release phage progeny (**Fig. 1D**). These results suggest that this defense system functions through regulated cell death or dormancy^22^ and provides population-level protection against phage T6. Homology-based searches in a database of ∼38,000 prokaryotic genomes revealed that this defense system is found in hundreds of bacteria and archaea belonging to diverse phyla (**Fig. 1E, Table S1**).

### Caspase cleaves EF-Tu during phage infection

TIR domains were described as NAD^+^-degrading effectors in diverse families of defense systems, including Pycsar, CBASS and prokaryotic argonautes, where their function is to deplete cells of NAD^+^ once triggered by phage infection^24–27^. Bacterial caspase-like proteases, on the other hand, were suggested to cleave bacterial gasdermins into active pore-forming effectors to induce cell death following infection^5^. For this reason, we initially hypothesized that *Ps*Caspase would cleave *Ps*TIR into an active NAD^+^-depleting form during infection. However, we did not detect any change in the molecular weight of *Ps*TIR during T6 infection (**Fig. S2A**). In addition, we observed only very mild reduction in NAD^+^ levels during infection of cells expressing the system, which contrasts with the >95% reduction typically observed in NAD^+^-depleting defense systems^13,26–30^ (**Fig. S2B**). These results suggest that *Ps*TIR is unlikely to be an NAD^+^-depleting effector activated by *Ps*Caspase.

In Thoeris defense systems, bacterial TIR domains have an alternative role that does not involve NAD^+^ depletion. In these systems, TIR domains generate signaling molecules that activate downstream effectors^10^. We therefore hypothesized that *Ps*TIR may produce a signaling molecule during infection, and that this signaling molecule would bind and activate the effector *Ps*Caspase. Under this hypothesis, *Ps*Caspase is expected to cleave cellular or phage target proteins and halt phage infection.

To explore possible target proteins of *Ps*Caspase, we chemically labeled NH_2_ groups in proteins present in lysates of T6-infected cells and subjected the labeled proteins to mass spectrometry (MS). This technique enables the identification of new protein N-termini specifically found in cells expressing the defense system as compared to control cells. A preliminary MS analysis following free amine tagging identified multiple candidate *Ps*Caspase targets, including the elongation factor Tu (EF-Tu), a highly abundant cellular protein essential for protein translation, which was previously shown to be the target of the Lit protease^31^ (**Fig. S3**). To verify that EF-Tu is a target of *Ps*Caspase, we co-expressed the TIR-caspase system with a C-terminally tagged copy of EF-Tu in an *E. coli* K-12 strain in which *lit* was deleted, and monitored EF-Tu cleavage during phage infection by western blot (**Fig. 2A**). Two EF-Tu cleavage products were visible in cells expressing the defense system following T6 infection (**Fig. 2B**). These cleavage products were absent when the catalytic cysteine in *Ps*Caspase or the catalytic glutamate in *Ps*TIR were mutated to alanine, showing that EF-Tu cleavage requires both caspase and TIR activities (**Fig. 2B**). The molecular weight of cleavage products and the exploratory MS data suggested that EF-Tu is cleaved after two specific arginine residues (R45 and R59) (**Fig. 2C**), which was confirmed by the loss of EF-Tu cleavage upon mutation of these two residues into glutamines (**Fig. 2B**). Taken together, our results suggest that *Ps*Caspase is specifically activated during phage infection and cleaves EF-Tu downstream of arginine residues at positions 45 and 59.

**Figure 2.**
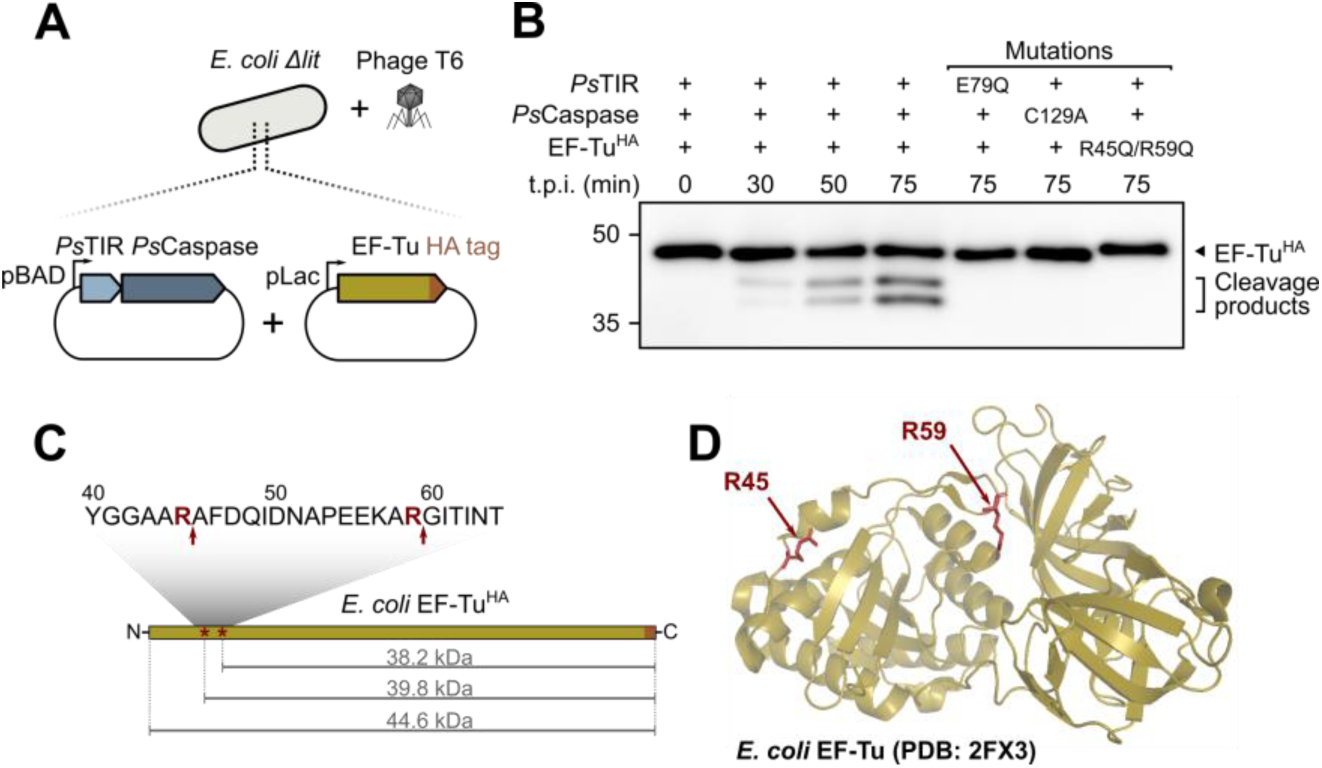
EF-Tu is a target of *Ps*Caspase during phage T6 infection. (**A**) *E. coli* K-12 MG1655Δ*lit* cells co-expressing the *Pseudomonas sp.* 1-7 defense system and an HA-tagged copy of EF-Tu were infected with phage T6. (**B**) Western blot analysis of infected cells co-expressing wild-type or mutated *Ps*TIR and *Ps*Caspase, and wild-type or mutated HA-tagged EF-Tu. t.p.i. : time post-infection. (**C,D**) Location of cleavage sites in the EF-Tu protein sequence (C) and structure (D) are indicated with red arrows.

### Caspase activity is triggered by a TIR-derived signaling molecule

We then investigated whether, as in the Thoeris family of defense systems, *Ps*TIR produces a signaling molecule that activates *Ps*Caspase during infection. To do so, we collected lysates from cells expressing *Ps*TIR only and filtered these lysates with a 3 kDa cutoff to retain small metabolites. We then tested the ability of these metabolite extracts to activate EF-Tu cleavage by *Ps*Caspase (**Fig. 3A**). EF-Tu cleavage was visible when lysates from cells expressing both *Ps*Caspase and tagged EF-Tu were incubated with metabolite extracts derived from infected *Ps*TIR-expressing cells (**Fig. 3B**). In contrast, EF-Tu cleavage was absent when the metabolites were extracted from infected cells expressing RFP or *Ps*TIR^E79Q^ instead of *Ps*TIR, or from uninfected *Ps*TIR-expressing cells (**Fig. 3B**). These results suggest that *Ps*TIR produces a *Ps*Caspase-activating signaling molecule during infection.

**Figure 3.**
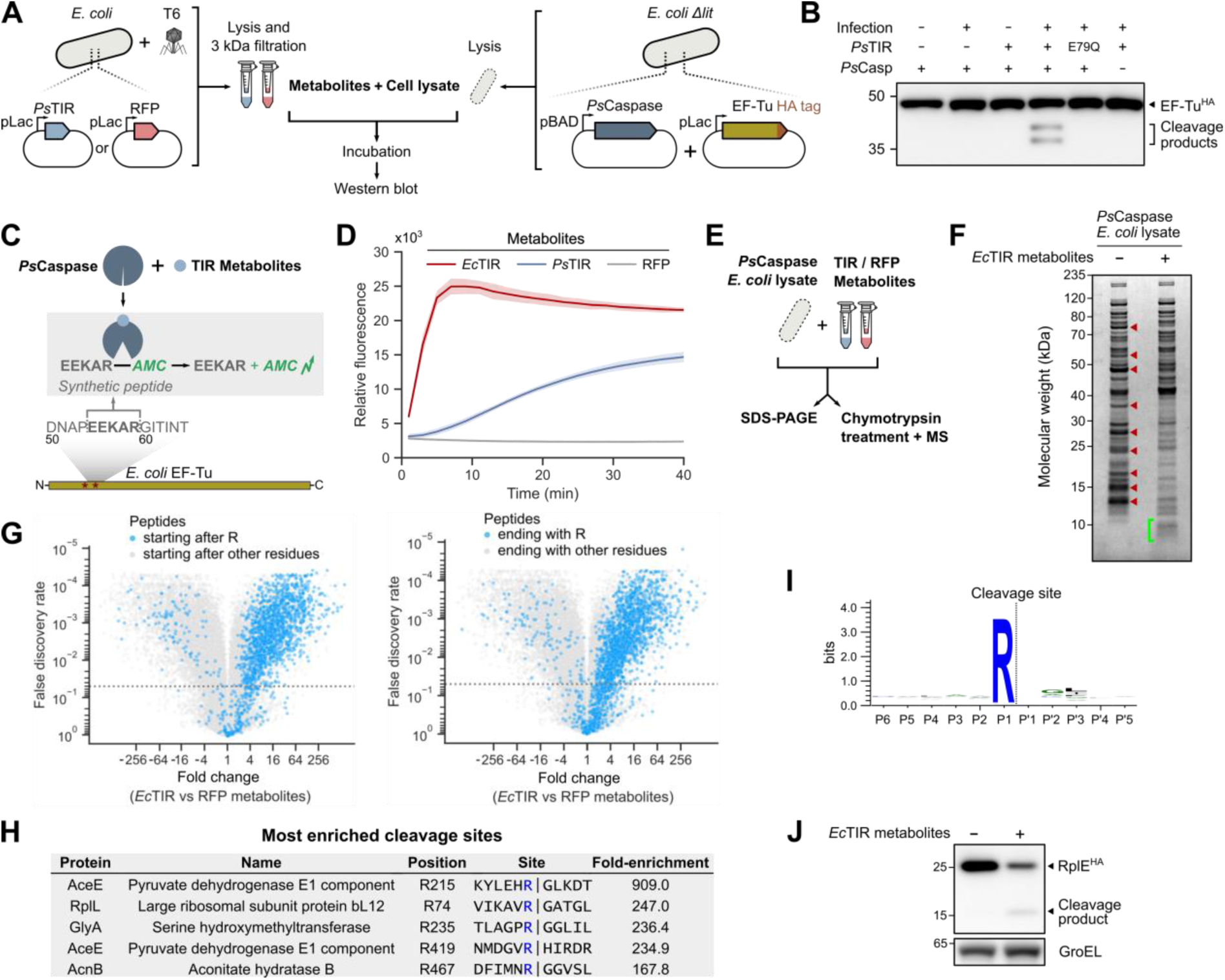
A TIR-derived signaling molecule triggers *Ps*Caspase arginine-specific proteolytic activity. (**A**) Experimental setup to test for *Ps*Caspase activation by small molecules. Lysates of T6-infected cells expressing *Ps*TIR or RFP were passed through 3 kDa filters to collect small molecules. Separately, *E. coli* K-12 MG1655Δ*lit* cells co-expressing *Ps*Caspase and a C-terminally HA-tagged copy of EF-Tu were lysed and incubated with the small molecules collected from infected *Ps*TIR- or RFP-expressing cells. Reactions were resolved by SDS-PAGE and western blot. (**B**) Western blot analysis shows EF-Tu cleavage only in the presence of metabolites derived from infected *Ps*TIR-expressing cells. (**C**) Schematic representation of the *Ps*Caspase reporter assay. A peptide comprising the residues preceding one of the cleavage sites in EF-Tu (EEKAR, positions 55-59) fused to 7-amino-4-methylcoumarin (AMC) was synthesized. The synthetic peptide was incubated with lysates from *Ps*Caspase-expressing *E. coli* K-12 MG1655Δ*lit* cells and small molecules collected from infected TIR- or RFP-expressing cells. *Ps*Caspase activation leads to peptide cleavage downstream of the arginine residue, thereby releasing the free fluorophore. (**D**) *Ps*Caspase reporter assay with small molecules collected from infected *Ec*TIR-, *Ps*TIR- or RFP-expressing cells. Curves show the mean of three replicates with the standard deviation shown as a shaded area. (**E**) Schematic of the *in vitro* cleavage assays, using lysates from *E. coli* K-12 MG1655Δ*lit* cells that express *Ps*Caspase, incubated with small molecules collected from T6-infected cells that express either *Ec*TIR or RFP. (**F**) SDS-PAGE of *in vitro* reactions followed by Coomassie staining. Red arrows indicate protein bands that are depleted in the presence of *Ec*TIR metabolites. Green bracket indicates a smear of low molecular weight fragments that are enriched in the presence of *Ec*TIR metabolites. (**G**) Volcano plots showing differential abundance of peptides in *Ps*Caspase cell lysates following incubation with *Ec*TIR-derived small molecules or with small molecules from control RFP-expressing cells. Peptides starting directly after an arginine residue (left) or ending with an arginine residue (right) are marked in blue. (**H**) Peptides enriched >5-fold in the *Ec*TIR sample, and depleted >2-fold when incubated with RFP vs *Ec*TIR metabolites were selected for motif analysis (n=83 peptides from 70 proteins). Shown are the five strongest hits. Fold-enrichment relates to peptides starting at the position after arginine. (**I**) Sequence logo of the 83 cleavage sites. (**J**) Cleavage of C-terminally HA-tagged RplE from cell lysates in the presence of *Ec*TIR metabolites as shown by western blot. GroEL is shown as a loading control.

To develop a more streamlined assay for *Ps*Caspase activation, we used a synthetic five amino acid peptide comprising the residues preceding the target cleavage site in EF-Tu (position 55-59, EEKAR) fused to 7-amino-4-methylcoumarin (AMC). In this assay, which is similar to assays previously used to study human caspases^32^ and caspase-like proteins in bacteria^8^, peptide cleavage downstream the arginine is expected to release free AMC and emit a fluorescence signal (**Fig. 3C**). *Ps*Caspase was able to cleave the synthetic EEKAR-AMC peptide when incubated with metabolites derived from infected cells expressing *Ps*TIR, but not when incubated with metabolites extracted from infected RFP-expressing control cells (**Fig. 3D**). *Ps*Caspase activity was even more pronounced upon incubation with metabolites derived from infected cells expressing the TIR protein from the *E. coli* 328 TIR-caspase system (*Ec*TIR) (**Fig. 3D**), suggesting that in the experimental conditions used here, *Ec*TIR produces higher amounts of signaling molecule than *Ps*TIR. Together, these results demonstrate that the caspase-like protein is activated by a small molecule specifically produced by the TIR-domain protein during phage infection. We therefore name this defense system type IV Thoeris.

### *Ps*Caspase is a promiscuous arginine-specific protease

We then investigated the substrate specificity of the caspase-like protein using synthetic peptides with mutations in the wild-type EF-Tu sequence (EEKAR). Activated *Ps*Caspase was able to cleave EE<u>A</u>AR-AMC and EEK<u>L</u>R-AMC peptides, but not an EEKA<u>A</u>-AMC peptide, indicating that the arginine residue at the P1 position is strictly required for cleavage, but other residues are not (**Fig. S4**). These results suggest that *Ps*Caspase does not specifically recognize EF-Tu but might rather be a promiscuous protease.

To systematically search for additional cellular targets of *Ps*Caspase beyond EF-Tu, we incubated lysates of *Ps*Caspase-expressing *E. coli* cells with metabolites extracted from infected *Ec*TIR-expressing cells (**Fig. 3E**). SDS-PAGE analysis revealed the loss of multiple abundant proteins in the presence of *Ec*TIR-derived metabolites, combined with the presence of a smear of low-molecular weight protein products, suggesting massive *Ps*Caspase-dependent protein degradation (**Fig. 3F**). We subjected these lysates to chymotrypsin digestion and mass spectrometry analysis to identify cleavage sites in a systematic manner. This analysis revealed that peptides starting directly downstream of arginine, or ending with arginine, were strongly enriched upon incubation with *Ec*TIR-derived metabolites (**Fig. 3G**), indicating that cleavage occurs downstream of arginine residues. Cleavage events were observed in multiple essential proteins of *E. coli* (**Fig. 3H**), with a strict requirement for an arginine in the P1 position and a slight preference for small nonpolar residues surrounding the cleavage site (**Fig. 3I**). We confirmed the cleavage of one of these target proteins, ribosomal protein RplE, by western blot using cell lysates incubated with *Ec*TIR-derived metabolites (**Fig. 3J**). Altogether, our results show that *Ps*Caspase is a promiscuous arginine-specific protease that cleaves multiple target proteins once activated by the *Ps*TIR-derived signaling molecule.

### Type IV Thoeris produces the new signaling molecule N7-cADPR

We next examined the properties of the *Ps*Caspase-activating signaling molecule produced by type IV Thoeris. For this, we sought to take advantage of previously described phage proteins that sequester specific Thoeris-derived signaling molecules as a means to inhibit bacterial defense. Thoeris anti-defense 1 (Tad1) from phage SBSphiJ7 and Tad2 from phage SPO1 are known to sequester the 1”-3’ gcADPR signaling molecule produced by type I Thoeris^12,33^, and Tad2 from *Myroides odoratus* (ModTad2) sequesters His-ADPR molecules and inhibits type II Thoeris^15^. However, co-expression of Tad1, Tad2 or ModTad2 with type IV Thoeris did not inhibit the defensive activity of the system **(Fig. 4A**), suggesting that type IV Thoeris does not utilize the same molecule as types I or II Thoeris. Furthermore, incubation of metabolites derived from *Ec*TIR-expressing infected cells with purified Tad proteins did not affect the ability of these metabolites to activate *Ps*Caspase in our reporter assay, suggesting that Tad proteins are unable to sequester the signaling molecule of type IV Thoeris (**Fig. S5A**). Finally, purified 1”-3’ gcADPR, 1”-2’ gcADPR, ADPR and cyclic-ADPR molecules were unable to activate *Ps*Caspase (**Fig. 4B**), and conversely, *Ec*TIR-derived metabolites were unable to activate the ThsA NADase effector of type I Thoeris^13^ *in vitro* (**Fig. S5B**). Together, these results indicate that the signaling molecule of type IV Thoeris is distinct from that of type I and type II Thoeris.

**Figure 4.**
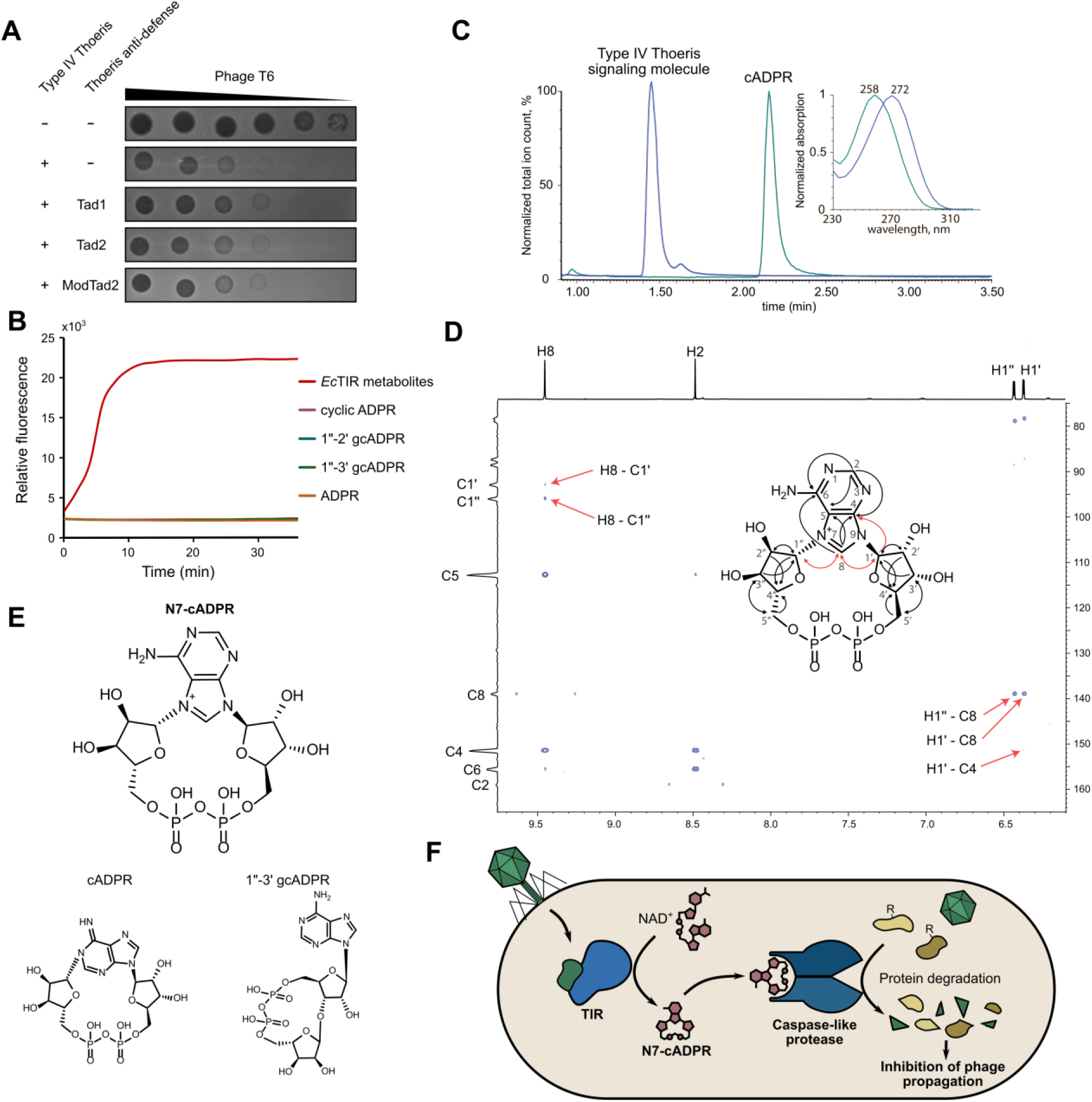
Type IV Thoeris produces the immune signaling molecule N7-cADPR. (**A**) Plaque assay of phage T6 on cells co-expressing the type IV Thoeris system from *Pseudomonas* sp. 1-7 with RFP control, Tad1 from phage SBSphiJ7^12^, Tad2 from phage SPO1^33^, or Tad2 from *M. odoratus* (ModTad2)^15^. None of the anti-defense proteins inhibits type IV Thoeris defense. (**B**) *Ps*Caspase reporter assays using metabolites derived from *Ec*TIR-expressing cells following phage infection, or using 100 µM of pure cyclic ADPR, 1”-2’ gcADPR,1”-3’ gcADPR or ADPR. Only *Ec*TIR-derived metabolites activate *Ps*Caspase. (**C**) Chromatograms and UV-spectra (inset) of the type IV Thoeris signaling molecule (blue) and canonical cADPR (N1-cADPR, green). Both molecules have m/z = 540.0543 in negative ionization mode. (**D**) Correlations observed for N7-cADPR in the 2D ^1^H-^13^C HMBC NMR spectrum. Correlations between the adenine and the two ribose moieties are marked by red arrows both in the spectrum and on the molecule; other HBMC-derived correlations are shown by black arrows on the molecule. (**E**) Structure of N7-cADPR, compared to structures of canonical cADPR and 1”-3’ gcADPR. (**F**) A model for the type IV Thoeris mechanism of immunity.

To further examine the nature of the signaling molecule produced by type IV Thoeris, we used HPLC fractionation to purify the molecule from lysates of phage-infected *Ec*TIR-expressing cells, using the peptide-AMC cleavage assay as a reporter system to assess the activity of HPLC fractions (**Fig. S6**). Initial purification in phosphate buffer (pH=8.0) showed that the molecule rapidly lost activity in these conditions, and we therefore purified it in water (pH=7.0). Liquid chromatography coupled with mass spectrometry (LC-MS) showed that the molecule has the same mass as the canonical cyclic ADP-ribose (cADPR) (**Fig. 4C**) and MS-MS analysis showed that the fragmentation pattern of the molecule was indistinguishable from that of cADPR (**Fig. S6D**). However, the molecule had a different LC retention time and absorbed light at a different wavelength than cADPR, suggesting that the type IV Thoeris molecule is distinct from cADPR (**Fig. 4C**). Indeed, synthetic cADPR was unable to activate *Ps*Caspase (**Fig. 4B**).

We subjected the purified molecule to Nuclear Magnetic Resonance (NMR) analysis. Data from 1D ^1^H, 2D homonuclear COSY and TOCSY, and 2D heteronuclear ^1^H-^13^C HSQC and HMBC NMR experiments showed that the molecule comprises two ribose moieties and an adenine base, supporting the notion that the molecule is a variant of cADPR (**Fig. S7**). However, ribose carbon 1’’, which in the canonical cADPR is naturally found attached to N1 in the adenine ring, showed a 3-bond HMBC correlation to H8 in the adenine ring (**Fig. 4D**, inferred from the H8-C1’’ and H1’’-C8 correlations in the HMBC spectrum). Moreover, similar intensities were observed for peaks correlating position 8 on the adenine base and 1’ and 1’’ in the two ribose moieties (H1’-C8 and H1’’-C8, and H8-C1’ and H8-C1’’), indicating a symmetrical orientation of the two ribose moieties relative to carbon atom C8 of the adenine base (**Fig. 4D**). Given that ribose carbon 1’ was verified as attached to N9 in the adenine base (H1’-C4 correlation), our results indicate that ribose carbon 1’’ is attached to nitrogen atom N7 in the adenine base (**Fig. 4D**). Altogether, our MS and NMR experiments indicate that the molecule that activates *Ps*Caspase is ADP-cyclo[N7:1′′]-ribose (N7-cADPR), a molecule that, to our knowledge, was not described before in natural systems (**Fig. 4E**). It was previously shown that N7-methylation of adenosine shifts the wavelength of the absorption maximum from 257 to 272 nm^34^, which is in line with the shift in absorption that we observe in N7-cADPR as compared to cADPR (**Fig. 4C**).

Together, our data provide a model for the mechanism of type IV Thoeris (**Fig. 4F**). Once infection is sensed in the cell, the TIR-domain protein utilizes NAD^+^ as a substrate and generates a cyclized ADPR molecule where the carbon previously connected to the nicotinamide ring becomes covalently attached to the N7 nitrogen atom in the adenine base. This N7-cADPR molecule specifically activates the caspase-like effector of the defense system which then indiscriminately cleaves cellular and potentially phage proteins after arginine residues, causing massive protein degradation in the cell and incapacitating the phage reproductive cycle (**Fig. 4F**).

## Discussion

In this study we discovered the molecule N7-cADPR and showed that it functions as an immune signaling molecule in bacteria. Past studies showed that immune signaling molecules tend to be shared between domains of life. For example, 2’3’ cyclic GMP-AMP functions as an immune signaling molecule in the human cGAS-STING pathway^35^, as well as in defense systems of the CBASS family^36^. Similarly, 3’3’ cyclic UMP-AMP, originally discovered as a signaling molecule in bacterial CBASS systems^37^, was later shown to be produced by animal immune proteins in response to dsRNA stimulation^38^. TIR-derived immune signaling molecules also have parallels between bacteria, animals and plants. For example, 1”-3’ gcADPR, produced by type I Thoeris, was shown to also be produced by the plant immune protein BdTIR^18^ and the human TIR-domain protein SARM1^39^, although the biological roles of this molecule in plants and human are currently unclear. It is therefore to be anticipated that the new molecule discovered here, N7-cADPR, could also be found to participate in immune signaling in multicellular eukaryotes in future studies.

Two types of Thoeris systems were previously studied in detail, and a third type (type III Thoeris) was suggested recently based on an operon architecture that contains TIR domains^16^. In type I Thoeris, the effector protein ThsA harbors a SLOG domain that binds the signaling molecule gcADPR^14^, and in type II Thoeris, the signaling molecule His-ADPR is perceived by a Macro domain found in the effector protein. In type IV Thoeris, the caspase-like protease effector encodes a C-terminal domain of unknown function, and we hypothesize that this domain is responsible for binding N7-cADPR to trigger the proteolytic activity of the effector (**Fig. S8A**). In support of this hypothesis, structural modeling suggests that *Ps*Caspase forms a dimer, in which the C-terminal domains are brought in close proximity to form a putative N7-cADPR-binding pocket (**Fig. S8B**). Future studies will be necessary to examine the mechanism of *Ps*Caspase activation.

Human caspases can function both as initiators of cell death, where they activate specific death-promoting proteins by proteolytic cleavage^1^, or as executioners of cell death, where they cleave hundreds of proteins to promote apoptosis^3^. Our findings strengthen the understanding that caspase-like proteins have similar roles in bacterial immunity. Bacterial caspase-like proteases were previously shown to cleave bacterial gasdermins into pore-forming effectors that execute a form of pyroptotic-like cell death^5^, paralleling inflammatory caspases in human^1,2^. In type IV Thoeris immunity, the role of the caspase-like protease is analogous to executioner caspases, as it directly executes cell death by cleaving many cellular target proteins, likely shutting off essential processes in the cell and preventing viral replication. Recent work has shown that some type III CRISPR-Cas systems also co-opted caspase-like proteases as a part of their immune mechanism^6–8^. In particular, a type III-B CRISPR-Cas system employs a cascade of proteolytic events, in which CRISPR-Cas signaling activates a SAVED-CHAT protease to specifically process *PCaspase*, a caspase-like promiscuous protease that cleaves multiple proteins possibly downstream of arginine residues^8^. Our results point to a common molecular function of *PCaspase* and the type IV Thoeris caspase as arginine-specific promiscuous proteases responsible for executing cell death or dormancy. Caspase-like proteases were predicted to be associated with other families of defense systems such as CBASS^40^ and Avs^41^, but their roles in these systems are currently unknown. It is possible that these proteases mediate bacterial defense through a mechanism similar to *Ps*Caspase. Our results show that proteases of the caspase family are ancient modules in the toolbox of immune-related cell death machineries shared by bacteria, fungi^42^, plants^43^, and humans^1^.

## Methods

### Bacterial strains and phages

*E. coli* NEB 5-alpha (New England Biolabs) was used as a cloning strain. *E. coli* K-12 MG1655 was used for phage-related experiments. The *lit* deletion was transferred to *E. coli* K-12 MG1655 from the Keio collection^44^ (clone ECK1125) using P1 transduction. *E. coli* BL21(DE3) was used to overexpress *Ec*TIR and *Ps*TIR to produce the type IV Thoeris signaling molecule. Unless stated otherwise, cells were grown in MMB (LB + 0.1 mM MnCl_2_ + 5 mM MgCl_2_, with or without 1.5% agar) with the appropriate antibiotics: ampicillin (100 mg/mL), chloramphenicol (30 mg/mL) or kanamycin (50 mg/mL).

### Plasmid construction

Type IV Thoeris systems from *E. coli* 328 (NCBI RefSeq assembly GCA_000503435.1) and *Pseudomonas sp.* 1-7 (NCBI RefSeq assembly GCA_000742775.1) were synthesized by Genscript into the pBAD vector^45^. *Ec*TIR and *Ps*TIR were cloned into the pBbA6c vector^46^ (NEBuilder HiFi DNA Assembly) C-terminally HA-tagged *E. coli tufA* (EF-Tu) and *rplE*, as well as Tad1 from phage SBSphiJ7^12^, Tad2 from phage SPO1^33^ and Tad2 from *M. odoratus* (ModTad2)^15^ were cloned by Gibson assembly into the pBbS6c vector^46^ (Addgene plasmid # 35290). Point mutations were introduced using the KLD Enzyme mix (New England Biolabs) and were verified by whole-plasmid sequencing (Plasmidsaurus). Cloned sequences are provided as Table S2.

### Plaque assays

Phage T6 was amplified from single plaques in liquid cultures of *E. coli* K-12 MG1655 at 37°C in MMB medium until culture collapse. Cultures were centrifuged (4,000 g x 10 min) and the supernatants were passed through 0.2 µm filters. Plaque assays were performed as previously described^47^. *E. coli* K-12 MG1655 carrying the pBAD plasmid encoding the type IV Thoeris system from *E. coli* 328 or *Pseudomonas sp.* 1-7 or a control GFP protein were grown overnight at 37°C in MMB medium supplemented with ampicillin. Then, 300 mL of each culture was mixed with 30 mL of molten MMB + 0.5% agar supplemented with 0.2% arabinose. The mixture was poured on 12 x 12 cm plates and left to dry for 1 h. Tenfold dilutions of phage T6 were prepared in MMB and 10 mL of each dilution was dropped onto the plates. Plates were incubated overnight at 37°C and plaque-forming units (PFUs) were counted the next day. To test Thoeris anti-defense proteins, plaque assays were performed in cells carrying the pBAD plasmid encoding the type IV Thoeris system from *Pseudomonas sp.* 1-7 as well as the pBbS6c vector^46^ encoding RFP, Tad1 from phage SBSphiJ7 ^12^, Tad2 from phage SPO1^33^ or Tad2 from *M. odoratus* (ModTad2)^15^. Plaque assays were conducted as described above in the presence of 0.2% arabinose and 500 µM IPTG.

### Infection assays in liquid cultures

Overnight cultures of *E. coli* K-12 MG1655 carrying the pBAD plasmid encoding the type IV Thoeris system from *Pseudomonas sp.* 1-7 or a control GFP protein were diluted 100-fold in MMB-ampicillin with 0.2% arabinose and grown to an optical density at 600nm (OD600) of 0.4. Then, 190 µL of cultures were transferred to a 96-well plate and mixed with 10 µL of MMB or phage T6 stock to reach an MOI of 3 or 0.03. Growth was followed by OD600 measurement every 5 min on a Tecan Infinite200 plate reader at 37°C. Propagation of phage T6 was measured by infecting the same exponential cultures at OD600 of 0.4 with phage T6 at an MOI of 0.1 (Fig. 1D). The infected cultures were sampled at the indicated time points, spun down at 14,000 *g* for 1 min, and the free phage titer was assessed in the supernatant by plaque assays on *E. coli* K-12 MG1655 as described above.

### Exploratory identification of free amines in *E. coli* proteins

Overnight cultures of *E. coli* K-12 MG1655 carrying the pBAD plasmid encoding the type IV Thoeris system from *Pseudomonas sp.* 1-7 or a control GFP protein were diluted 100-fold in MMB-ampicillin with 0.2% arabinose and grown to an OD600 of 0.4. Cells were then infected with T6 at an MOI of 3 and 5 mL of cells were collected 40 min post infection. Bacterial cell pellets were lysed by addition of 50 μL of 8 M urea buffer. Lysates were sonicated for 10 cycles (30s on/ 30s off; Diagenode). Protein concentration was measured using the BCA assay (Thermo Scientific, USA). From each sample, 15 ug of total proteins were reduced with 5 mM dithiothreitol (Sigma) for 1 h at room temperature and alkylated with 10 mM iodoacetamide (Sigma) in the dark for 45 min at room temperature. Samples were diluted to 2 M urea with 50 mM TEAB. Free amines (lysines and protein N-termini) were then labeled with 0.8 mg TMT10plex (Thermo Fisher) according to the manufacturer’s instructions, including quenching of unreacted TMT tags. Proteins were then subjected to digestion with trypsin (Promega; Madison, WI, USA) overnight at 37°C at 50:1 protein:trypsin ratio, followed by a second trypsin digestion (50:1 protein:trypsin ratio) for 4 h. The digestions were stopped by addition of trifluroacetic acid (1% final concentration). Following digestion, peptides were desalted using Oasis HLB, μElution format (Waters, Milford, MA, USA). The samples were vacuum dried and stored in -20°C until further analysis.

ULC/MS-grade solvents were used for all chromatographic steps. Dry digested samples were dissolved in 97:3% H_2_O/acetonitrile + 0.1% formic acid. Each sample was loaded using split-less nano-Ultra Performance Liquid Chromatography (10 kpsi nanoAcquity; Waters, Milford, MA, USA). The mobile phase was: A) H_2_O + 0.1% formic acid and B) acetonitrile + 0.1% formic acid. Desalting of the samples was performed online using a reversed-phase Symmetry C18 trapping column (180 µm internal diameter, 20 mm length, 5 µm particle size; Waters). The peptides were then separated using a T3 HSS nano-column (75 µm internal diameter, 250 mm length, 1.8 µm particle size; Waters) at 0.35 µL/min. Peptides were eluted from the column into the mass spectrometer using the following gradient: 4% to 25% mobile phase B in 155 min, 25% to 90% mobile phase B in 5 min, maintained at 90% for 5 min and then back to initial conditions. The nanoUPLC was coupled online through a nanoESI emitter (10 μm tip; New Objective; Woburn, MA, USA) to a quadrupole orbitrap mass spectrometer (Q-Exactive HFX, Thermo Scientific) using a FlexIon nanospray apparatus (Proxeon).

Data was acquired in data dependent acquisition (DDA) mode, using a Top10 method. MS1 resolution was set to 120,000 (at 200 m/z), mass range of 375-1650 m/z, AGC of 10^6^ and maximum injection time was set to 60 ms. MS2 resolution was set to 15,000, quadrupole isolation 1.7 m/z, AGC of 10^5^, dynamic exclusion of 45 s and maximum injection time of 60 ms.

The data was analyzed using Proteome Discoverer v2.4 (Thermo Scientific). The raw data were searched against a protein database containing the recombinant protein sequences, the *E. coli* K-12 and T6 bacteriophage protein databases as downloaded from Uniprot, and a list of common lab contaminants. The search was done with two search algorithms, SequestHT and MS Amanda, allowing identification of semi-tryptic peptides. The following modifications were defined for the search: i) Fixed modification: cysteine carbamidomethylation; ii) Variable modifications: methionine oxidation, and TMT label on lysines and protein/peptide N-terminus.

### Western blotting

Overnight cultures of *E. coli* K-12 MG1655Δ*lit* cells carrying (i) the pBAD plasmid encoding the wild-type or mutated (*Ps*TIR^E89Q^ or *Ps*Caspase^C129A^) type IV Thoeris system from *Pseudomonas sp.* 1-7 or a control GFP protein, and (ii) the pBbA6c vector encoding the C-terminally HA-tagged wild-type or mutated (R45Q/R59Q) EF-Tu, were diluted 100-fold in MMB supplemented with ampicillin, chloramphenicol, 0.2 % arabinose and 100 µM IPTG and were grown at 37°C with shaking. When OD600 reached 0.5, 250 µL of each culture were sampled, centrifuged (14,000*g* for 1 min) and pellets were frozen. The remaining cultures were infected with phage T6 at an MOI of 3, and the sampling procedure was repeated 30, 50 and 75 min post-infection. All frozen pellets were resuspended in 200 µL of 1X Bolt LDS Sample Buffer supplemented with 50 mM DTT. Samples were incubated at 95°C for 3 min and 10 µL were separated by 4–12% Bis-Tris SDS-PAGE (Thermo Scientific) in 1x MOPS buffer (Thermo Scientific). Gels were transferred to a nitrocellulose membrane (Invitrogen #LC2001) for 1 hour at 20 V in 1x transfer buffer (Thermo Scientific). Membranes were blocked in 5% milk in TBS-T buffer for 30 min with shaking and probed with a rabbit anti-HA antibody (Sigma Aldrich #H6908; 1:5,000 dilution). Visualization was performed using HRP-conjugated goat anti-rabbit secondary antibody (Thermo Scientific #31460; 1:10,000 dilution) and incubation with ECL solution (Merck Millipore). Where appropriate, membranes were stripped using Restore PLUS Western Blot Stripping Buffer (Thermo Scientific) and probed with primary rabbit anti-GroEL antibody (Sigma Aldrich #G6532, 1:10,000 dilution) as loading control.

### Preparation of metabolite fractions

Overnight cultures of *E. coli* K-12 MG1655 carrying the pBbA6c vector^46^ encoding RFP, *Ec*TIR or *Ps*TIR were diluted 100-fold in 200 mL of MMB supplemented with chloramphenicol and 500 µM IPTG. When OD600 reached 0.5, 100 mL were collected by centrifugation (4,000 *g* for 8 min) and pellets were frozen. Remaining cultures were infected with phage T6 at an MOI of 3 and collected 50 min post-infection. All pellets were resuspended in 1 mL of 100 mM sodium acetate (pH 5.5), transferred to a FastPrep Lysing Matrix B 2 ml tube (MP Biomedicals) and lysed using a FastPrep bead beater for 40 s at 6 m.s^-1^. Samples were centrifuged (12,000 *g* for 10 min) and supernatants were passed through 3 kDa Amicon Ultra-0.5 Centrifugal Filter Units (Merck Millipore, catalogue no. UFC500396) for 45 min at 12,000 *g* at 4°C. Metabolite fractions were stored at -80°C until further use.

### Peptide cleavage reporter assays

To produce *Ps*Caspase cell lysates, an overnight culture of *E. coli* K-12 MG1655Δ*lit* carrying the pBAD plasmid encoding *Ps*Caspase was diluted 100-fold in 50 mL of MMB supplemented with ampicillin and 0.2% arabinose. When OD600 reached 0.8, cells were collected by centrifugation (4,000 *g* for 8 min) and the pellet was resuspended in 500 µL of 100 mM sodium phosphate buffer (pH 7.4), resulting in *Ps*Caspase cell lysates. Peptide cleavage reporter assays were prepared by combining 1 µL of *Ps*Caspase cell lysates with 2 µL of 100 µM synthetic 7-amino-4-methylcoumarin (AMC)-conjugated peptides (Genscript) and 5 µL of 100 mM sodium phosphate buffer (pH 7.4) in a black 384-well plate (Corning 4511). Reactions were started by adding 2 µL of *Ec*TIR, *Ps*TIR or RFP metabolites. AMC fluorescence (excitation/emission wavelength: 341/441 nm) was measured on a Tecan Infinite M200 plate reader every 2 min relative to a pure solution of 10 µM AMC.

### ThsA NADase assay

ThsA NADase assays were performed as previously described^13^: briefly, in a volume of 45 µL, purified ThsA protein^13^ (final concentration of 100 nM) was mixed with RFP or *Ec*TIR metabolites derived from infected cells, or with pure 1”-3’-gcADPR (final concentration of 5 µM) in a black 96-well half area plates (Corning, 3694). Five microliters of 5 mM nicotinamide 1,N^6^-ethenoadenine dinucleotide (εNAD, Sigma, N2630) solution was added to each well immediately before the beginning of measurements and mixed by pipetting. Fluorescence was measured in a Tecan Infinite M200 plate reader every minute at 25°C at 300 nm excitation wavelength and 410 nm emission wavelength.

### Large scale production of the signaling molecule

Overnight cultures of *E. coli* K-12 MG1655 cells carrying *Ec*TIR were diluted 100-fold in 6 L of MMB supplemented with chloramphenicol and 500 µM IPTG. When OD600 reached 0.5, IPTG concentration was increased to 1 mM (final) and cells were grown for 1 extra hour, then the culture was infected with phage T6 at an MOI of 3 and collected after 1 h. Cells were resuspended in 2:1 (mL/g) of water and mixed with 1 volume (g/g) of glass beads (Sigma G4649) and vortexed at high speed for 1 h at 4°C, then lysozyme was added to 1mg/ml and the lysate was incubated for 1 h on ice. Lysate was frozen in liquid nitrogen and thawed to room temperature, the supernatant was then separated by centrifugation at 4000 *g* for 15 min. The pellet was dissolved in water again and extraction was repeated. The whole procedure was repeated 5 times. Mixed supernatant was passed through Amicon® Ultra-15 Centrifugal Filter Unit 3 kDa at 4,000 g at 4°C, flowthrough was passed through bond Elut C18 cartridge (10 g, 60 mL, 120 μm) by gravity force and flowthrough was lyophilized and dissolved in 2 mL of 0.03% Formic acid.

### HPLC

HPLC of the obtained fraction was performed using Agilent 1260 and chromatography Polar RP column Synergi by three separation steps. The following protocol was used for all runs – 5 min of mobile phase A 100%, 2 min 75% A and 25% B, 3 min 50% A and 50% B, 5 min 20% A and 80% B and 5 min 100 % A, 3 mL/min flow rate. For the first and third separation steps, mobile phase A was 0.03% formic acid and B was 0.03% formic acid in 20% methanol. For the second separation step, A was 20mM potassium phosphate pH 6 and B was 20mM potassium phosphate pH 6 in 20% methanol. During the first separation, half-minute fractions were collected and tested by peptide cleavage reporter assays. The active fraction was lyophilized, dissolved in 20mM potassium phosphate pH 6 and separated. HPLC peak with N7-cADPR characteristic spectrum (273 nm maximum) was collected, lyophilized and re-cleaned again in 0.03% formic acid to remove potassium phosphate.

### LC-MS

The sample solutions were analyzed by UPLC–HRMS without dilution. The analyses were carried out on a Waters SYNAPT-XS Q-Tof mass spectrometer (Manchester, UK) with an electrospray ionization (ESI) source. Lock spray was acquired with Leucine Encephalin (*m*/*z*=554.2615 in negative ionization mode and *m*/*z*=556.2771 in positive ionization mode) at a concentration of 200 ng/mL and a flow rate of 10 μL/min once every 10 s for 1 s period to ensure mass accuracy. Waters MassLynx v4.2 software was used for data acquisition and data processing. The spectra of N1-cADPR and N7-cADPR were recorded in the negative ionization mode within a mass range from 50 to 600 *m*/*z*. The parameters were set as follows: capillary voltage at 1.5 kV, cone gas flow at 50 L/h, source temperature at 140°C, and cone voltage at 20 V(−). The desolvation temperature was set at 600°C, and the desolvation gas (N2) flow rate was set at 800 l/h. Trap collision energy ramped from 20 to 50V and transfer collision energy ramped from 10 to 30V. The analytes were separated using Waters Premier Acquity UPLC system. The aqueous mobile phase consisted of 0.5% of formic acid (Fisher Scientific A117-50). The isocratic elution was achieved with Waters Acquity Premier HSS T3 Column, 1.8 μm, 2.1 × 100 mm at 0.25 mL/min flow rate, 35°C.

### NMR Spectroscopy

Dry compound was dissolved in 60 μL ^2^H_2_O (Tzamal D-Chem Laboratories Ltd), adding 0.01% 3-propionic-2,2,3,3-d4 acid sodium salt (TMSP, that was used as an internal chemical shift reference for ^1^H and ^13^C spectra) and transferred to a 1.7-mm NMR micro test tube. NMR spectra were recorded on a Bruker AVANCE NEO-600 NMR spectrometer equipped with a 5 mm TCI-xyz CryoProbe (using a special NMR spinner turbine to hold the 1.7 mm test tube in a 5 mm probe). All spectra were acquired at 279 K. All ^1^H and ^13^C chemical shift assignments were based on different 1D and 2D NMR techniques, all using pulsed field gradient selection. Chemical shifts (δ) are reported in parts per million (ppm) and splitting patterns are abbreviated as follows: s, singlet; d, doublet; t, triplet; m, multiplet.

The structure of N7-cADPR was determined by 1D ^1^H NMR spectrum, 2D homonuclear ^1^H-^1^H COSY and TOCSY spectra, and 2D ^1^H-^13^C heteronuclear HSQC and HMBC spectra. 1D ^1^H NMR spectra were acquired using 32 k data points and a recycling delay of 2 s. 2D homonuclear ^1^H-^1^H COSY and TOCSY spectra were acquired using 8192 (t2) x 400-800 (t1) data points. The 2D TOCSY spectrum was recorded using a dipsi2 mixing scheme of 120 ms. 2D heteronuclear ^1^H-^13^C HSQC and HMBC spectra were recorded using 4096 (t2) x 400 (t1) data points. Multiplicity-edited HSQC enables differentiating between methyl and methine groups that give rise to positive correlation, *vs* methylene groups that appear as negative peaks. The evolution delay of the heteronuclear multiple-bond correlation (HMBC) experiment was set to observe long range couplings of JH,C = 8 Hz.

Chemical shift assignment was based on combined information derived from one- and two-dimensional NMR spectra: 1D ^1^H, 2D homonuclear COSY and TOCSY correlations (**Fig. S7A-D** and **Table S4**), as well as 2D heteronuclear ^1^H-^13^C HSQC and HMBC spectra (**Fig. S7-E-F** and **Table S4**). Identification of the ribose moieties was based on 2D homonuclear COSY and TOCSY spectra, used to delineate connectivity between ^1^H signals within the individual sugar units (**Fig. S7B-D** and **Table S4**). This assignment was further supported by heteronuclear correlations (**Fig. S7E-F & Table S4**). Assignment of the adenine ^1^H and ^13^C shifts was based on heteronuclear multiple-bound HMBC correlations, observed between ^1^H and ^13^C that are 2-3 bonds apart (**Figs. 4D, S7E-F** and **Table S4**). Observation of weak HMBC cross peaks between H2-C5 and H8-C6 further supports the assignment, attributing these to weak, 4-bond correlation (**Figs. 4D & S7F-G** and **Table S4**). The slow chemical exchange observed for the ^1^H signal at 9.45 ppm (data not shown) further supports its assignment as H8, known to reveal slow exchange rates^48^. At the absence of protons which are 3-bonds apart, the cyclization details could only be revealed by long-range ^1^H-^13^C HMBC correlations. Two strong cross peaks between the adenine and the two ribose moieties, H1’-C8 and H2”-C8, in addition to correlations between H8-C1’, H8-C1” and H1’-C4 observed in the HMBC spectrum (**Fig. S7H**, marked by red), indicate that the enzyme catalyzes the formation of C1”(ribose)-N7 and C1’(ribose)-N9. ^1^H-^13^C HMBC-derived correlations are shown in **Fig. 4D**: correlations connecting the adenine with the two ribose moieties are marked by red arrows; other long-range HMBC correlations are shown by black arrows. The down-field shift of H8 (9.73 ppm) that becomes deshielded further supports this cyclization: a similar effect was observed for N7-substituted guanosine (N7-guanosome)^49^.

### *In vitro* cleavage assays and mass spectrometry

*Ps*Caspase cell lysates and metabolites derived from infected cells expressing *Ec*TIR or RFP (control) were produced as described above. Cleavage assays were performed by mixing 5µL of *Ps*Caspase lysates with 2 µL RFP or *Ec*TIR metabolites (collected after T6 infection) and 8 µL of 100 mM sodium phosphate (pH 7.4). Reactions were performed in triplicates and incubated at 37°C for 60 min. One microliter of each reaction was separated on a 4–12% Bis-Tris SDS-PAGE (Thermo Scientific) in 1x MOPS buffer (Thermo Scientific). For mass spectrometry analysis, *E. coli* cells were subjected to urea lysis, protein reduction and alkylation, and proteolytic digestion as described above in *“Exploratory identification of free amines in E. coli proteins“,* with the exception of omitting TMT-labeling, and using chymotrypsin (Sigma, St. Louis MO, USA) instead of trypsin. Chromatographic procedures were also as described above with the exception of elution gradient starting from 4% to 30% mobile phase B in 105 min, 30% to 90% B in 10 min, maintained at 90% B for 7 min and then back to initial conditions. The nanoUPLC was coupled online through a nanoESI emitter (10 μm tip; New Objective; Woburn, MA, USA) to a quadrupole orbitrap mass spectrometer (Exploris 480, Thermo Scientific) using a FlexIon nanospray apparatus (Proxeon).

Data was acquired in data dependent acquisition (DDA) mode, using a 2-seconds cycle time method. MS1 resolution was set to 120,000 (at 200 m/z), mass range of 380-1500 m/z, normalized AGC target of 200% and custom maximum injection time. MS2 resolution was set to 15,000, quadrupole isolation 1.4 m/z, normalized AGC target of 75%, dynamic exclusion of 40 s and maximum injection set to auto.

Raw data were processed with the MetaMorpheus v1.0.2 platform^50^. Peptide sequences were searched against a protein database comprising the *E. coli* proteome downloaded from Uniprot, the recombinant *Ps*Caspase protein, and a list of common lab contaminants. Parameters allowing for identification of semi-specific chymotryptic peptides were specified. Quantitative comparisons were calculated using Perseus v1.6.15.0^51^. Decoy hits were filtered out, and only proteins that were detected in at least two replicates of at least one experimental group were kept. Missing intensity values went through imputation (replaced by a random distribution of low values) for statistical purposes. For each protein, we recorded the amino acid positions where at least one detected peptide ends, or starts at the next position. For each of these positions, we calculated the median enrichment of peptides that either start at the next position, end at the position, or include the position (neither as first or last residue), in samples incubated with *Ec*TIR vs RFP metabolites (**Table S3**). Across all proteins, *Ps*Caspase cleavage sites were stringently selected as positions filling the three combined criteria: (i) >5-fold median enrichment of peptides starting at the next position, (ii) >5-fold median enrichment of peptides ending at the position, and (iii) >2-fold median depletion of peptides spanning the position. These criteria selected 83 cleavage sites disseminated in 70 proteins, 81 of which (97.6%) were found at arginine positions. These 83 sites were used to compute a sequence logo using WebLogo3^52^.

### Detection of type IV Thoeris in microbial genomes

The protein sequence of *Pseudomonas* sp.1-7 caspase-like protein (IMG gene ID 2600496627) was searched in a database of 3,895 finished prokaryotic genomes (3,781 bacterial and 114 archaeal)^29^ using the search function of MMseqs2 (release 12-113e3)^53^ with 3 iterations and otherwise default parameters. Homologs were aligned using Clustal-omega^54^ and the resulting multiple sequence alignment was used to build an HMM profile using the hmmbuild function of hmmer (version 3.3.2)^55^. In parallel, the HMM profiles of the TIR_2 (PF13676) and ThsB (PF08937) protein families were downloaded from the InterPro database^56^. MacSyFinder^57^ was then used to search the system in a database of 38,167 bacterial and archaeal genomes downloaded from the Integrated Microbial Genomes (IMG) database^58^ in October 2017: MacSyFinder was run with parameters --coverage-profile 0.1 --i-evalue-sel 0.1, requiring the caspase-like protein profile and either TIR_2 or ThsB HMM profiles to be detected, with up to one other gene between each system gene. The resulting detected systems were manually inspected for false positives. All detected systems are shown in **Table S1**.

## Acknowledgements

We acknowledge Corine Katina for preparing protein samples for mass spectrometry, and all members of the Sorek lab for constructive criticism on the manuscript. F.R. was supported by the Clore Foundation postdoctoral fellowship and by the Dean of Faculty fellowship from the Weizmann Institute of Science. I.O. was supported by the Ministry of Absorption New Immigrant program. Tali Scherf is the incumbent of the Monroy-Marks Research Fellow Chair. R.S. was supported, in part, by the European Research Council (grant ERC-AdG GA 101018520), the Israel Science Foundation (MAPATS grant 2720/22), the Deutsche Forschungsgemeinschaft (SPP 2330, grant 464312965), the Minerva Foundation with funding from the Federal German Ministry for Education and Research, the Ernest and Bonnie Beutler Research Program of Excellence in Genomic Medicine, a research grant from the Estate of Marjorie Plesset, the Institute for Environmental Sustainability (IES) and the Center for Immunotherapy at the Weizmann Institute of Science, and the Knell Family Center for Microbiology. M.I. and S. M. are supported by the Vera and John Schwartz Family Center for Metabolic Biology.

## Author contributions

F.R., I.O. and R.S. conceptualized the study. F.R. conducted phage infection experiments, bioinformatic detection of type IV Thoeris and EF-Tu cleavage experiments, and analyzed cleavage sites in mass spectrometry data. I.O. produced and purified the signaling molecule and analyzed analytical chemistry data. T.S. performed and analyzed the NMR experiments. A.H.F. performed LC-MS analyses of N7-cADPR and cADPR. A.L. performed cloning, G.A. devised the peptide-based reporter assay, S.S. tested Thoeris anti-defense proteins, M.I. and S.M. performed LC-MS experiments and A.S. performed protein mass spectroscopy analyses. F.R., R.S. and I.O. wrote the manuscript.

---

**Supplementary Figure 1.**
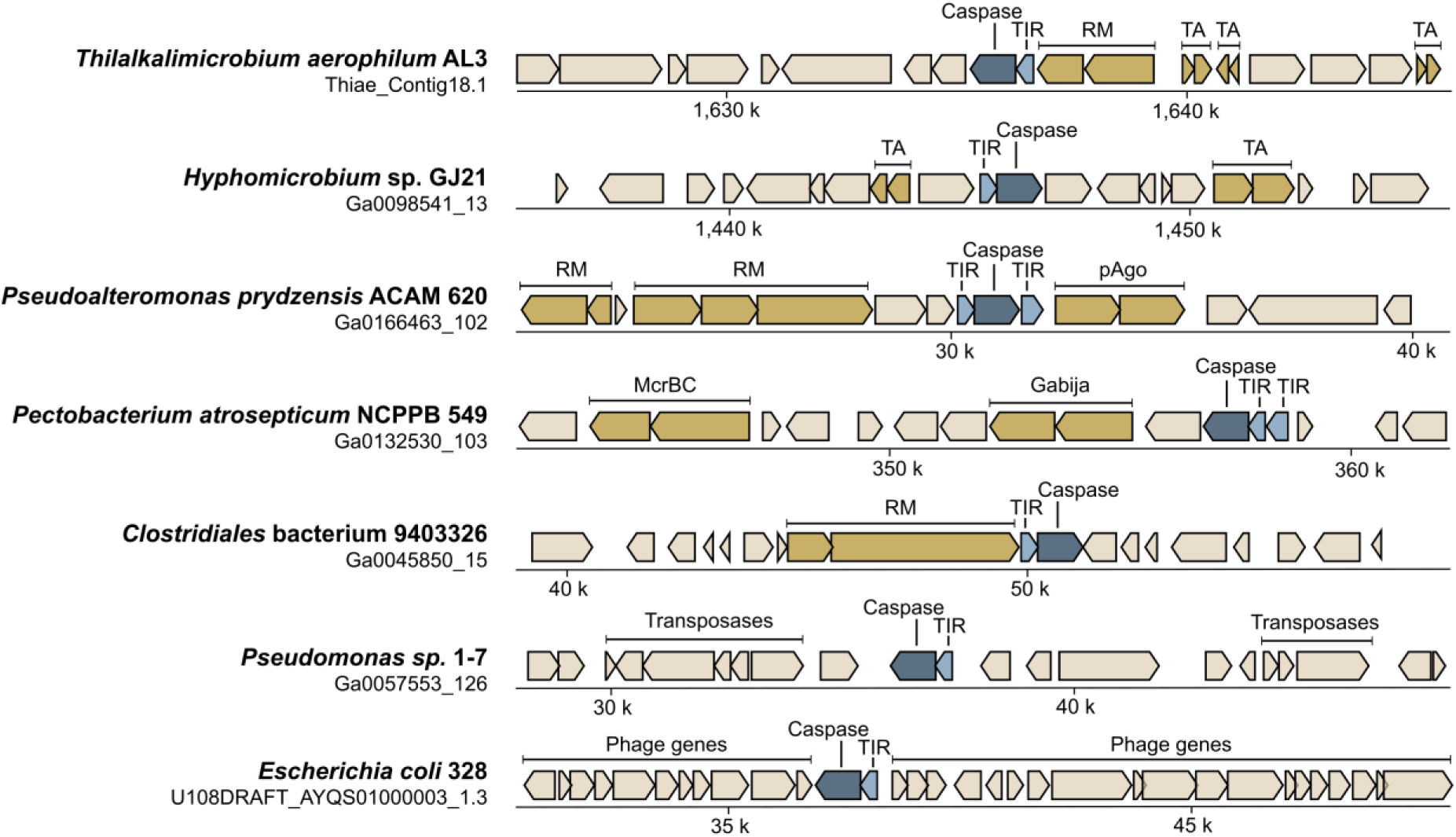
Operons encoding a TIR-domain protein and a caspase-like protease are frequently encoded near defense systems or on mobile genetic elements. Known defense systems are displayed in yellow. The IMG accession ID of bacterial contigs is displayed below genome names.

**Supplementary Figure 2.**
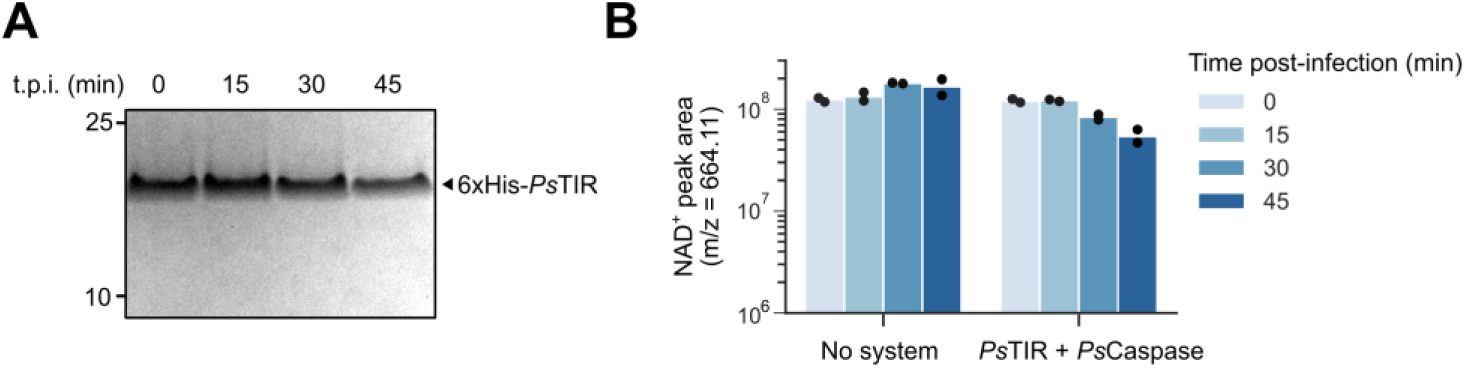
*Ps*TIR is unlikely to be a caspase-activated NAD^+^-degrading effector. (**A**) *E. coli* K-12 MG1655 cells expressing 6xHis-tagged *Ps*TIR and untagged *Ps*Caspase were infected with phage T6 at a MOI of 3. *Ps*TIR was pulled-down using Ni-NTA beads and samples were separated by 4-12% Bis-Tris SDS-PAGE followed by Coomassie staining. t.p.i: time post infection. (**B**) NAD^+^ levels in infected cells. *E. coli* K-12 MG1655 cells expressing *Ps*TIR and *Ps*Caspase, or a control GFP protein, were infected with phage T6 at a MOI of 3 and were collected at various time points. Metabolite fractions were extracted from lysates of collected cells and subjected to LC-MS analysis to record NAD^+^ levels.

**Supplementary Figure 3.**
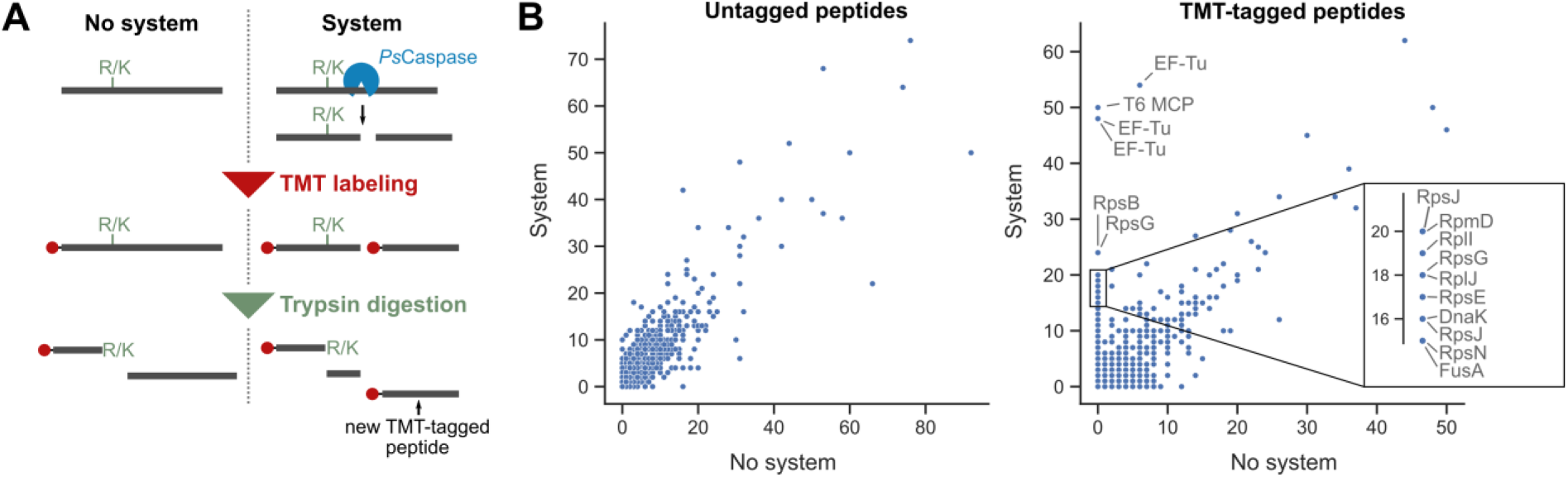
Exploratory protein mass spectrometry analysis identifies candidate targets for *Ps*Caspase. (**A**) Schematic representation of the experimental process. Protein lysates from infected cells encoding a control GFP or the defense system were labelled on free amines with tandem mass tags (TMT) and subsequently subjected to trypsin digestion. (**B**) Enrichment of specific TMT-tagged peptides in cells encoding the defense system. Each dot represents an amino acid position in a given protein. Shown is the number of untagged (left) or TMT-tagged (right) unique peptides recoded in lysates from T6-infected cells expressing GFP (x-axis) or the defense system (y-axis). Lysates were collected 40 minutes post infection by T6.

**Supplementary Figure 4.**
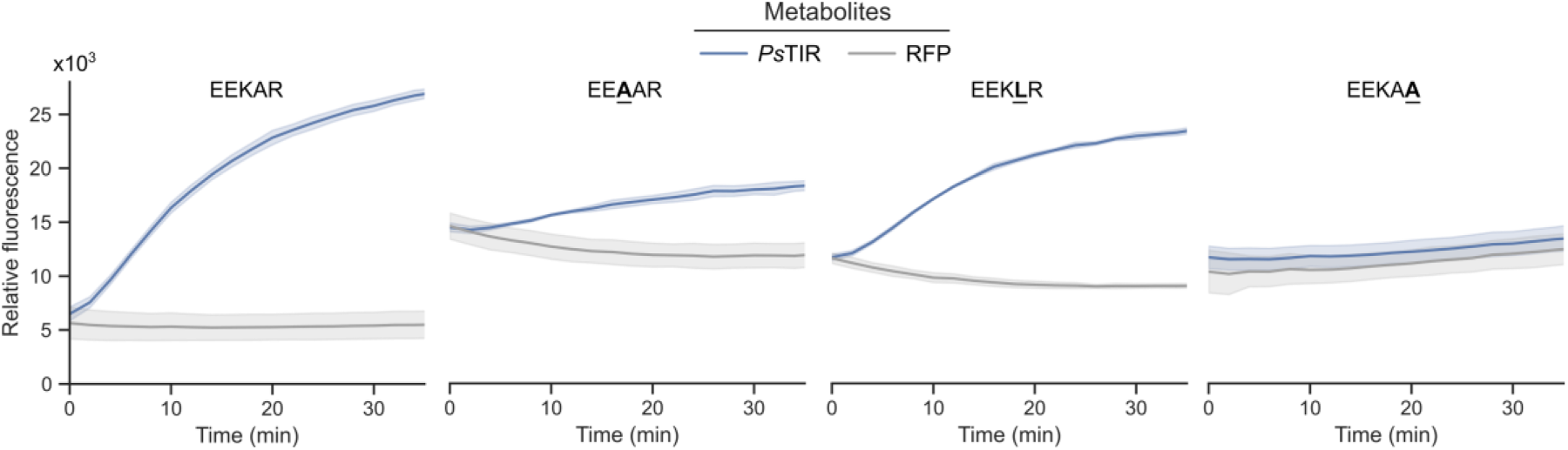
Cleavage of synthetic peptides by *Ps*Caspase. Cell lysates from *Ps*Caspase-encoding cells were incubated with metabolite fractions derived from T6-infected cells expressing *Ps*TIR or RFP (control) in the presence of synthetic peptides C-terminally fused to the fluorophore AMC. Peptide cleavage releases free AMC and emits a fluorescent signal (excitation/emission: 341/441 nm).

**Supplementary Figure 5.**
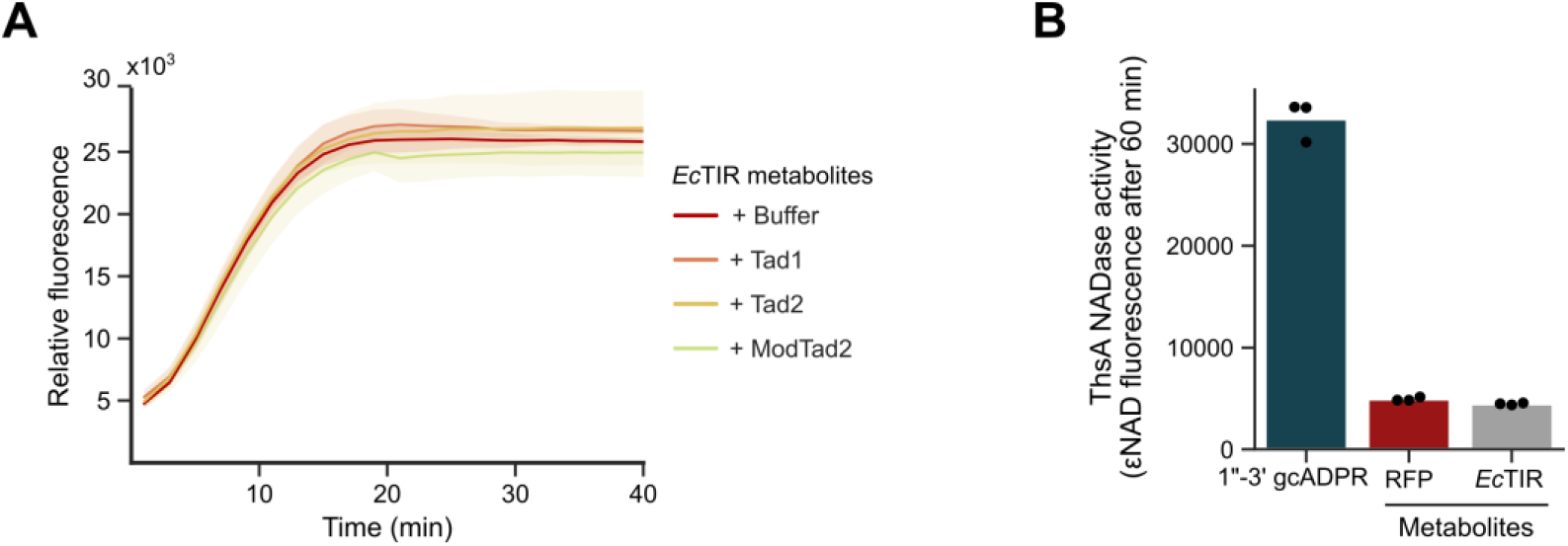
Type IV Thoeris signaling involves a molecule distinct from that of type I and type II Thoeris systems. (**A**) *Ps*Caspase reporter assay using metabolites derived from infected cells that express *Ec*TIR. These metabolites were pre-incubated with 4 µM of purified Tad1, Tad2 or ModTad2 for 15 minutes at room temperature. (**B**) ThsA NADase activation assay using metabolites derived from infected cells that express *Ec*TIR, RFP negative control or 1”-3’ gcADPR as a positive control^59^.

**Supplementary Figure 6.**
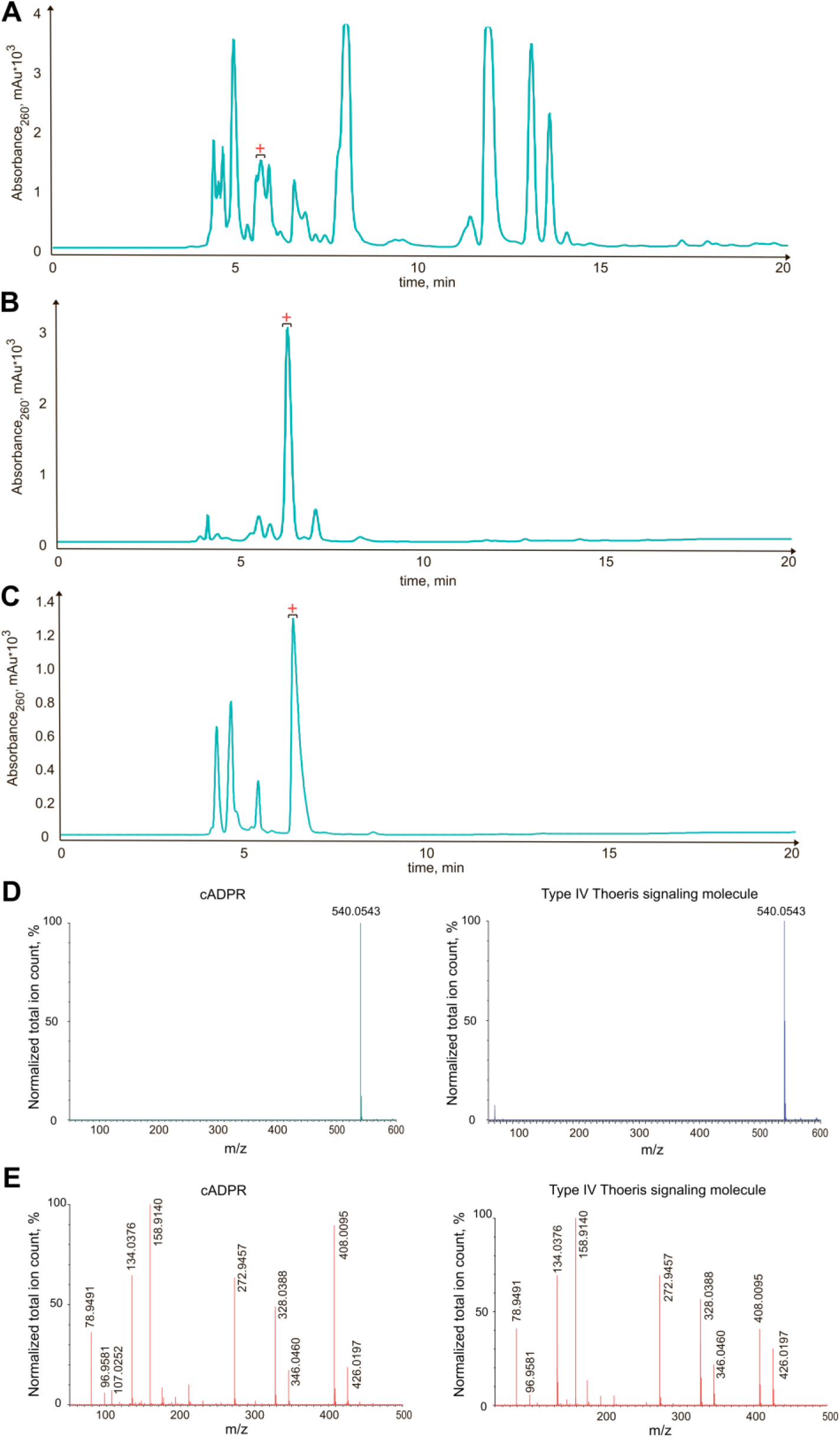
Chromatographic separation and purification of N7-cADPR and comparative MS-MS analysis of cADPR and N7-cADPR. Metabolite extract derived from T6-infected cells expressing *Ec*TIR were subjected to HPLC purification by three separation steps (A, B and C). **A.** Step 1: Mobile phase A, 0.03% formic acid; mobile phase B, 0.03% formic acid in 20% methanol. **B.** Step 2: Mobile phase A, 20mM potassium phosphate pH 6; mobile phase B, 20mM potassium phosphate pH 6 in 20% methanol. **C.** Step 3: Mobile phase, 0.03% formic acid; mobile phase B, 0.03% formic acid in 20% methanol. Active fractions collected for the next purification step are marked with a red plus sign (+). (**D,E**) MS analysis (**D**) and MS-MS analysis (**E**) of canonical cADPR (N1-cADPR) and N7-cADPR in negative ionization mode.

**Supplementary Figure 7.**
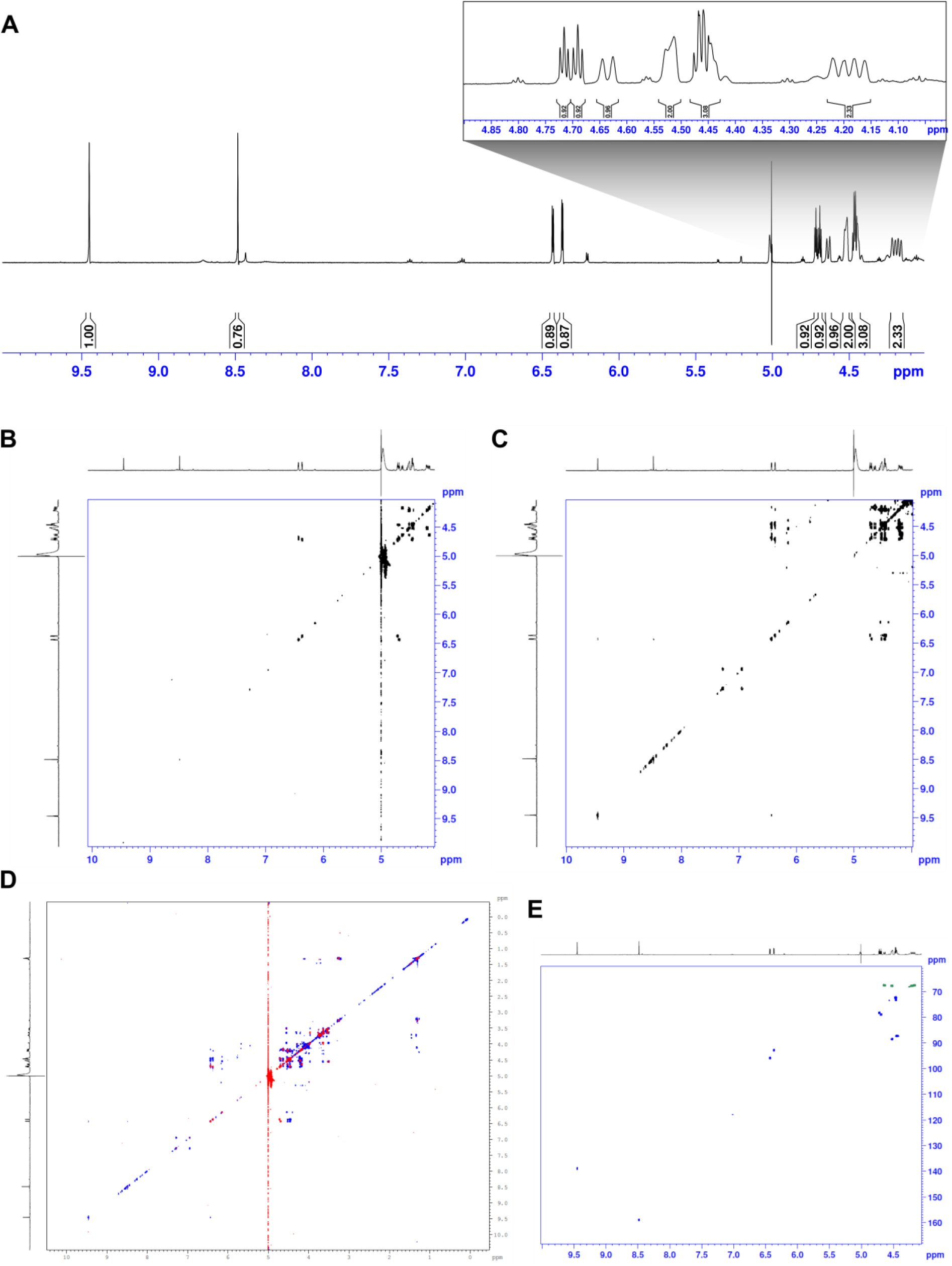
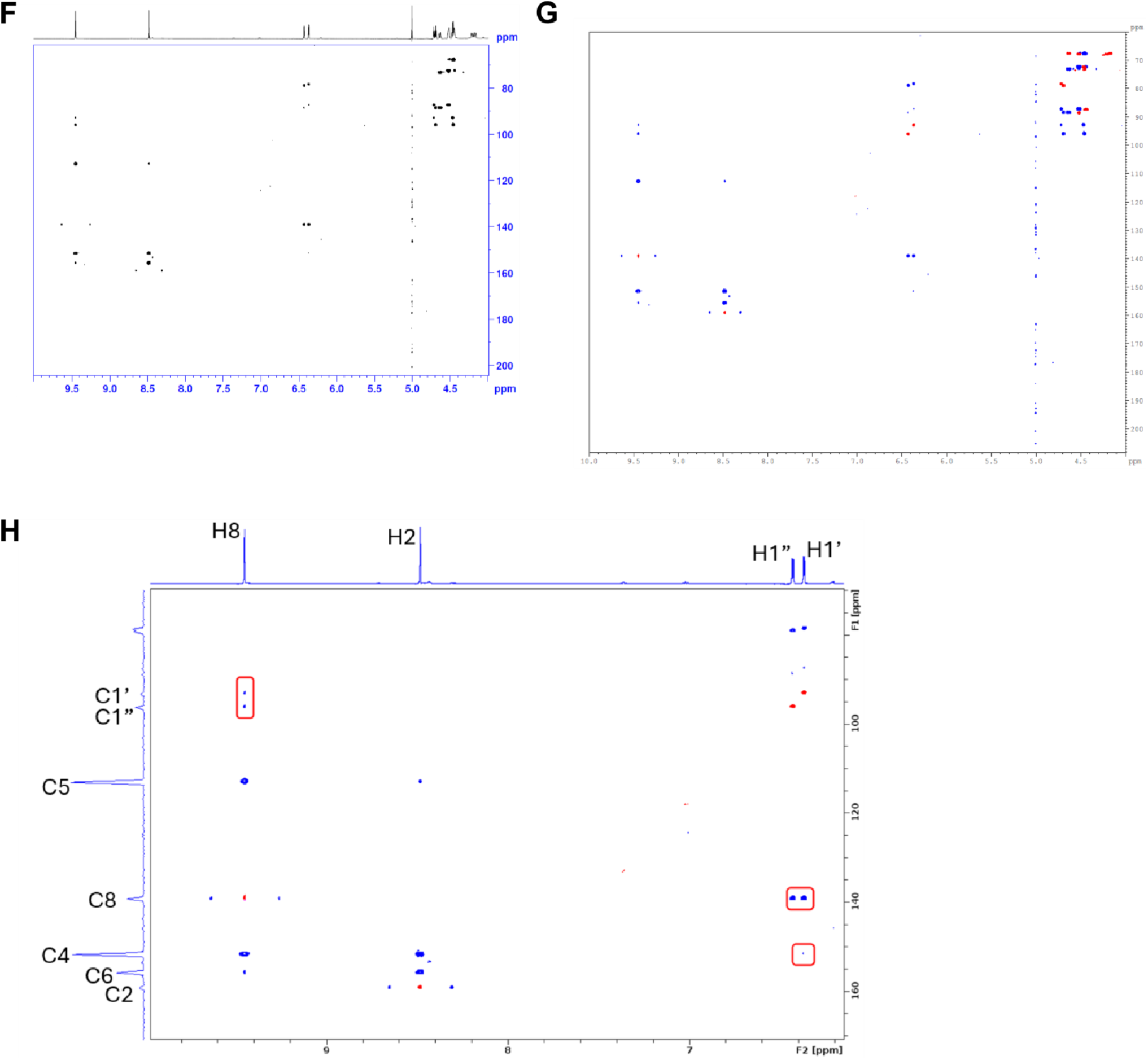
Nuclear Magnetic Resonance (NMR) analysis of the type IV Thoeris signaling molecule. (**A**) 1D ^1^H NMR spectrum of N7-cADPR (600 MHz, D2O), 279 K. (**B**) 2D ^1^H-^1^H COSY spectrum of N7-cADPR (600 MHz, D2O), 279 K. (**C**) 2D ^1^H-^1^H TOCSY spectrum of N7-cADPR (600 MHz, D2O), 279 K, using dipsi2 mixing scheme of 120 ms. (**D**) Superposition of the 2D ^1^H-^1^H TOCSY spectrum (blue), on the COSY spectrum (red). (**E**) 2D ^1^H-^13^C multiplicity-edited HSQC spectrum of N7-cADPR (600 MHz, D2O), 279 K. Methyl and methine groups are shown in blue, *vs* methylene groups in green. (**F**) 2D ^1^H-^13^C HMBC spectrum of N7-cADPR (600 MHz, D2O), 279 K. Long-range coupling was set to 8 Hz. Enlargement is shown as Fig. 4D. (**G**) Superposition of the 2D ^1^H-^13^C HMBC spectrum (blue), on the multiplicity-edited HSQC spectrum (red & purple). (**H**) Superposition of the 2D ^1^H-^13^C HMBC spectrum (blue), on the multiplicity-edited HSQC spectrum (red), revealing correlations of H2, H8, H1’ and H1” protons (enlargement of (**G**)). Correlations between the adenine and the two ribose moieties are marked in red.

**Supplementary Figure 8.**
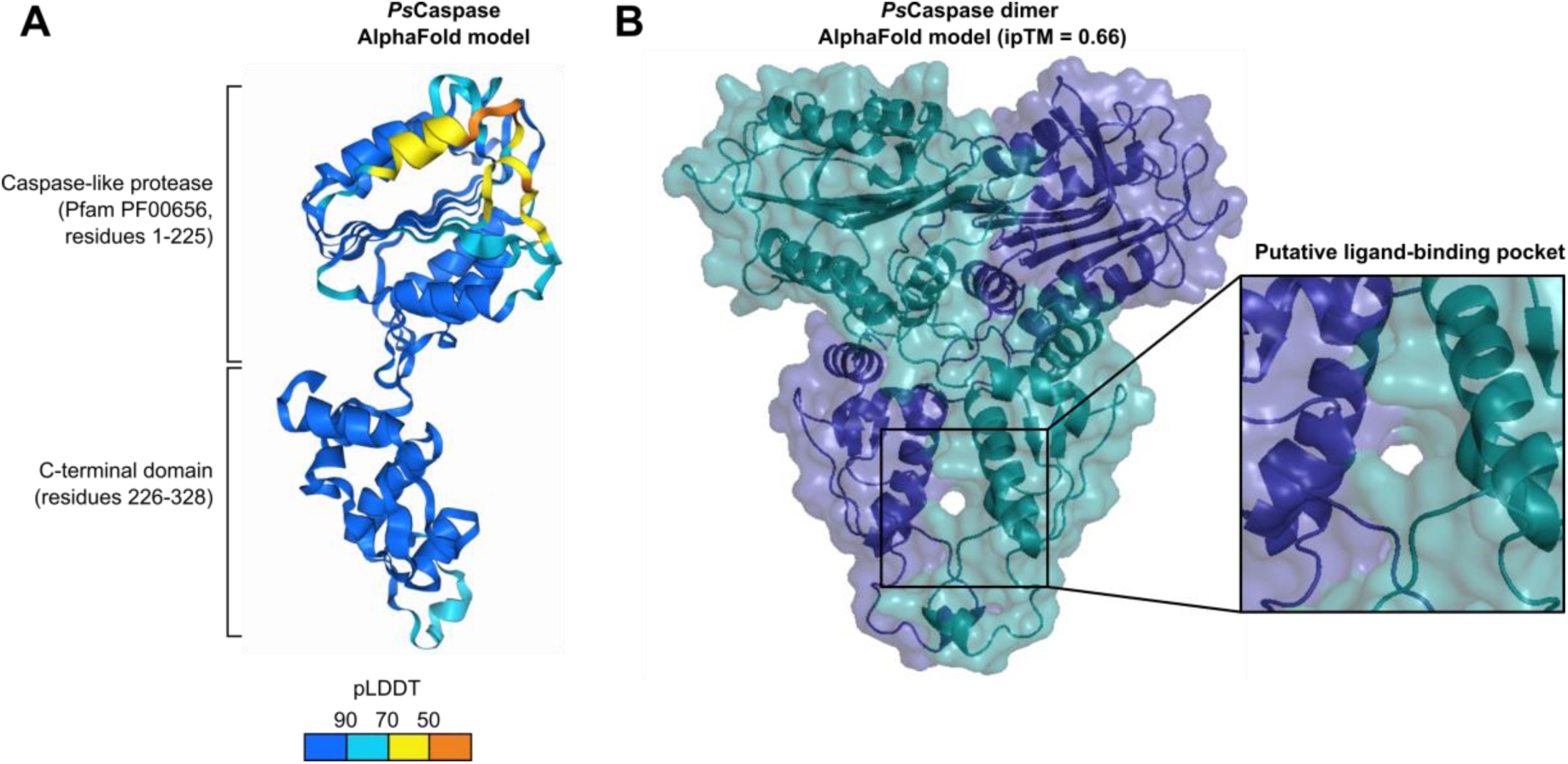
Structural modeling of *Ps*Caspase in monomeric and dimeric forms. (**A**) AlphaFold3^60^ model of the *Ps*Caspase monomer. Residues are colored by pLDDT score. (**B**) AlphaFold3^60^ model of the *Ps*Caspase dimer, colored by chain.

